# Co-immunization with pre-erythrocytic antigens alongside circumsporozoite protein can enhance sterile protection against *Plasmodium* sporozoite infection

**DOI:** 10.1101/2022.06.17.496580

**Authors:** Vladimir Vigdorovich, Hardik Patel, Alexander Watson, Andrew Raappana, Laura Reynolds, William Selman, Suzannah Beeman, Paul T. Edlefsen, Stefan H.I. Kappe, D. Noah Sather

## Abstract

Malaria-causing *Plasmodium* parasites have a complex life cycle and present numerous antigen targets that may contribute to protective immune responses. The currently recommended vaccine—RTS,S—functions by targeting the *P. falciparum* circumsporozoite protein (CSP), which is the most abundant surface protein of the sporozoite form responsible for initiating infection of the human host. Despite showing only moderate efficacy, RTS,S has established a strong foundation for the development of next-generation subunit vaccines. Our previous work characterizing the sporozoite surface proteome identified additional non-CSP antigens that may be useful as immunogens individually or in combination with CSP. In this study, we examined eight such antigens using the rodent malaria parasite *P. yoelii* as a model system. We demonstrate that despite conferring weak protection individually, co-immunizing each of several of these antigens alongside CSP, could significantly enhance the sterile protection achieved by CSP immunization alone. Thus, our work provides compelling evidence that a multi-antigen pre-erythrocytic vaccine approach may enhance protection compared to CSP-only vaccines. This lays the groundwork for further studies aimed at testing the identified antigen combinations in human vaccination trials that assess efficacy with controlled human malaria infection.

**Importance:** The currently approved malaria vaccine targets a single parasite protein (CSP) and only results in partial protection. We tested several additional vaccine targets in combination with CSP to identify those that could enhance protection from infection upon challenge in the mouse malaria model. In identifying several such enhancing vaccine targets, our work indicates that a multi-protein immunization approach may be a promising avenue to achieving higher levels of protection from infection. Our work identified several candidate leads for follow-up in the models relevant for human malaria, and provides an experimental framework for efficiently carrying out such screens for other combinations of vaccine targets.

## Introduction

Since the year 2000, the world has observed a steady decline in the global burden of malaria, but this trend has plateaued for the past six years, hinting at the insufficiency of current global efforts for malaria eradication. In 2020, the COVID-19 pandemic caused logistical interruptions in malaria prevention and treatment programs resulting in a significant increase in global malaria burden, with an estimated 241 million new cases and 627 thousand deaths, 77% of which were children under 5 years of age (1). It is clear that current interventions, including distribution of insecticide-treated bednets for mosquito control and the use of several classes of drugs for the treatment of infection, are susceptible to disruption due to numerous factors, such as political instability, resistance to insecticides, and emerging resistance to antimalarial drugs. Further, the slowing gains in eradication outcomes prior to the pandemic indicate that these efforts may not be sufficient to achieve eradication without the addition of new prevention measures. Thus, although significant progress in malaria control and patient survival following infection has taken place since the turn of this century, achieving the future eradication of malaria will likely rely on the ongoing work to develop highly effective malaria vaccines.

Malaria disease is caused by multiple species of obligate intracellular parasites belonging to the *Plasmodium* genus. The sporozoite forms of the parasite are inoculated into the skin of a mammalian host by an infected *Anopheles* mosquito. Sporozoites then traverse the skin tissues in search of a blood vessel that they may invade to be carried with the blood circulation to the liver. Once in the liver, the sporozoites exit the circulatory system and, using unknown means, choose hepatocytes to invade and establish infections. In the course of liver infection, each of the 10–1000 sporozoites (2) deposited by a mosquito bite can establish a liver stage, each producing 10,000 or more merozoite forms (3, 4) that exit the liver and initiate the blood-stage infection cycle in the host erythrocytes. Thus, targeting the relatively low-abundance sporozoite form of the parasite before the dramatic expansion of the parasite mass in the liver presents a critical opportunity for malaria vaccine design. Importantly, prevention of infection at this stage would stop both clinical disease and transmission.

The most advanced malaria vaccine—RTS,S/AS01—is based on a recombinant version of the circumsporozoite surface protein (CSP), which is highly abundant on the sporozoite surface. The WHO has recommended RTS,S/AS01 for use in young children because it has shown to decrease incidence of severe malaria by ~30% (1, 5, 6). A similar subunit vaccine—R21/Matrix-M —has recently undergone a smaller phase 2b trial and yielded high antibody titers with a higher vaccine efficacy, compared to RTS,S/AS01, against clinical malaria (7, 8). Multiple studies strongly associated the magnitude and quality of antibody response with protection elicited by these vaccines (7, 9–12). The R21/Matrix-M study also reported that although antibody titers wane within a year, they could be boosted to near-initial levels (7), while maintaining high efficacy (8). Thus, CSP-based subunit vaccines have established a foundation for the development of next-generation malaria vaccines that may be achievable by including additional antigens from the *Plasmodium* parasite.

Recent work has shown the association of protection with responses to non-CSP antigens (13). When applied in the vaccine context, whole sporozoite vaccine (WSV)-based approaches have shown that immunization with radiation- or chemically-attenuated sporozoites can induce antibody responses against a broad spectrum of *P. falciparum* antigens including CSP (14–16). Notably, the WSV-induced antibodies to non-CSP sporozoite antigens have been shown to inhibit sporozoite invasion of hepatocytes, implying that such antigens may elicit protective immune responses against pre-erythrocytic (PE) stage infection (17). Similar vaccine-elicited sporozoite inhibition was recently shown more directly in the *P. yoelii* (*Py*) rodent malaria model by active co-immunization with CSP alongside a panel of PE antigens normally expressed during liver stage development (18). However, protection in those experiments was assessed using an i.v. challenge, which, unfortunately, can miss protective mechanisms active in the skin (19).

Previously, we used mass spectrometry to identify surface-exposed proteins in different *Plasmodium* species (20–22). Among the hits were proteins that localize either to the sporozoite surface or those found in the invasion organelles (micronemes and rhoptries), ready to be released due to specific stimuli associated with the initiation of invasion. In addition, we showed that many components of the inner membrane complex could become exposed during the initial process of invasion and be accessible for recognition by antibodies (22). Recently, we demonstrated that passively transferred monoclonal antibodies (mAbs) raised against TRAP can enhance the protective effect of anti-CSP mAbs (23). In the present study we investigated whether active immunization with non-CSP sporozoite antigens could enhance the anti-CSP response-mediated protection using the rodent parasite model. To achieve this, we developed a platform for evaluating synergistic antigen combinations that could potentially provide guidance for human malaria vaccine development. We screened recombinant versions of eight *Py* sporozoite-expressed proteins as PE immunogens in pairwise combination with CSP to test their ability to enhance the CSP-mediated sterile protection in mice upon mosquito-bite challenge. Overall, we identified four antigens, which, when used in combination with CSP, enhanced sterile protection over CSP alone, providing new strategies for the future development of effective multi-antigen vaccines to prevent malaria infection.

## Results

### Single-antigen vaccination with recombinant Py sporozoite antigens elicits variable degrees of infection blocking despite strong antibody responses

To establish the potential efficacy of single-protein vaccine formulations against liver infection, we selected ectodomains of nine PE vaccine antigens expressed by *Py* sporozoites for recombinant production and purification alongside an unrelated control protein, HIV-1 Env gp120 (Table 1, Fig. 1A). Constructs encoding ectodomains of *Py* proteins: circumsporozoite protein (PyCSP), GPI-anchored micronemal antigen (PyGAMA), sporozoite surface protein 3 (PySSP3), thrombospondin-related anonymous protein (PyTRAP), thrombospondin-related sporozoite protein (PyTRSP), P36 (PyP36), P52 (PyP52), cell-traversal protein for ookinetes and sporozoites (PyCelTOS) and heat-shock protein 70-2 (PyHSP70-2) were engineered for production in HEK293 cells. Additionally, the PyCSP construct contained two point mutations to disrupt the N-linked glycosylation motif and prevent this post-translational modification. Groups of mice (n=3) were primed with 20 μg of individual *Py* sporozoite antigen in 20% Adjuplex adjuvant and 14 days later received an identical booster dose (Fig. 1B). Adjuplex is an experimental adjuvant that is a biodegradable matrix of carbomer homopolymer (Carbopol) and nano-liposomes derived from soy lecithin (24), and is known to stimulate robust antibody titers for recombinant protein vaccine formulations.

**Table 1.** Constructs.

| **<u>Construct name</u>** | **<u>Parasite localization</u>** | **<u>Mutations</u>** | **<u>UniProt reference</u>** | **<u>Domain(s)</u>** | **<u>Amino Acids</u>** |
| --- | --- | --- | --- | --- | --- |
| PyCSP | surface | S27A,T348A <sup>a</sup> | A0A077Y0S6 | Ectodomain | 21–362 |
| PyGAMA | micronemal | n/a | A0A077Y6C3 | Ectodomain | 22–603 |
| PySSP3 | surface | n/a | A0A077YBH7 | Ectodomain | 23–387 |
| PyTRAP | micronemal | n/a | A0A077YAH0 | Ectodomain | 23–752 |
| PyTRSP | surface | n/a | Q6IMC2 | Ectodomain | 21–145 |
| PyP52 | micronemal | n/a | A0A077YGW1 | Ectodomain | 24–458 |
| PyP36 | micronemal | n/a | A0A077YEG1 | Ectodomain | 71–356 |
| PyCeITOS | micronemal | n/a | Q6T944 | Ectodomain | 25–185 |
| PyHSP70-2 | ER, surface | n/a | A0A077Y5M2 | Ectodomain | 22–651 |
| HIV-1 Env gp120 | n/a | n/a | P19550 | gp120 | 30–492 |

<sup>a</sup> PyCSP mutations that disrupt the N-glycosylation motifs.

**Figure 1.**
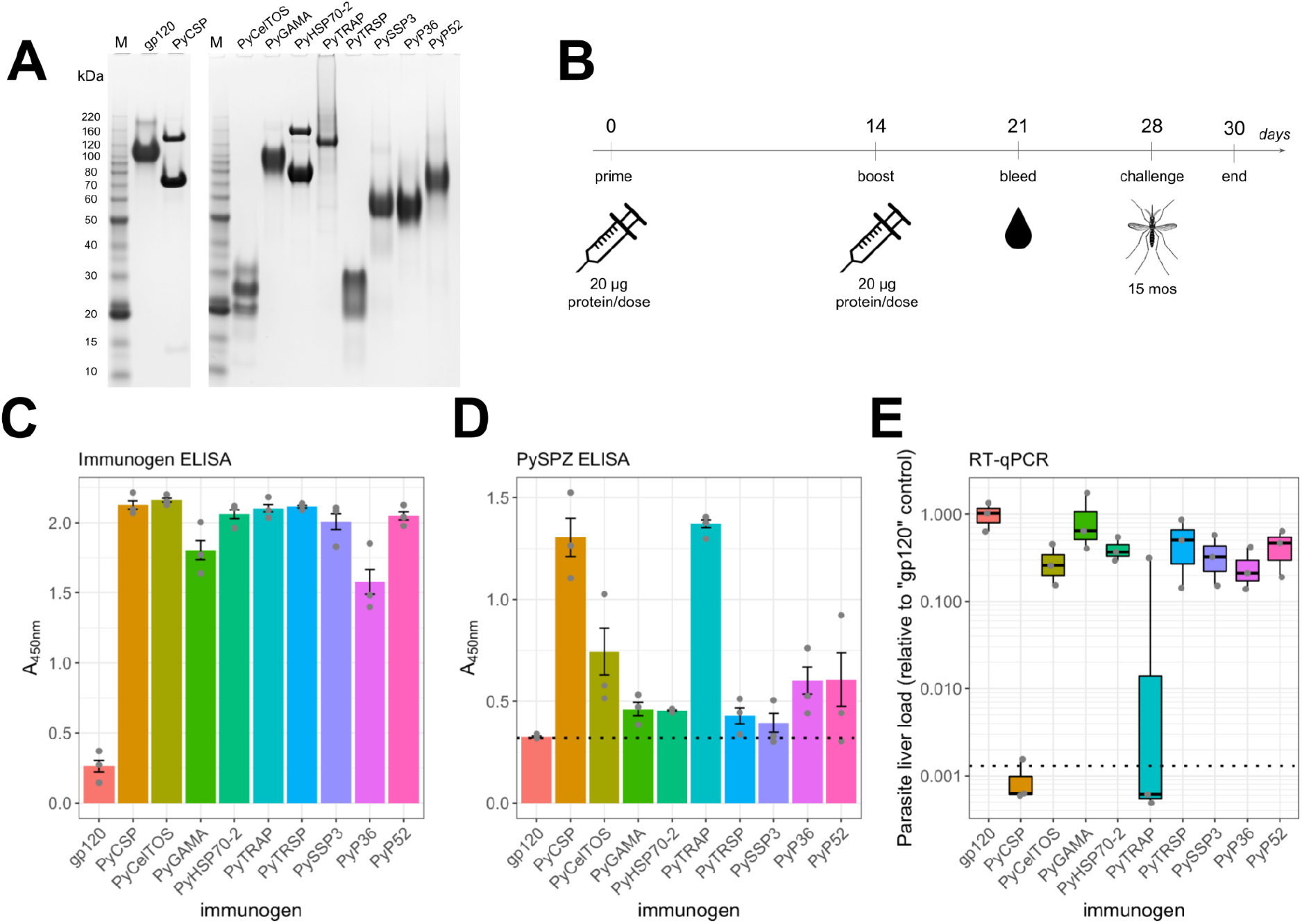
Immunization with recombinant Py PE antigens induces specific antibody responses but varying degrees of protection against infection in the liver. (**A**) SDS-PAGE analysis of the nine recombinant *Py* PE antigens used in immunizations and a control protein, HIV-1 Env gp120 (‘gp120’). (**B**) The immunization-challenge strategy schematic. Ten groups of mice (*n*=3) were immunized twice with each of PE antigens (in 20% Adjuplex) 14 days apart, followed by blood sample collection on day 21 and challenged with bites from 15 *Py*-infected mosquitoes on day 28. Parasite liver load was determined by RT-qPCR after harvesting the livers on day 30 (45h post-challenge). (**C, D**) The bar graphs show total antibody reactivity in 1:50-diluted “Day-21” plasma to purified PE antigens (**C**, each immunization group was tested against its corresponding antigen) or to whole sporozoites (**D**) by ELISA, with bar heights corresponding to the mean signal and the error bars representing ±SEM. In (**D**), the empirical background level for the sporozoite samples is indicated by a dotted line corresponding to reactivity against the control (‘gp120’) protein. (**E**) The parasite liver load (*Py* 18S rRNA) quantified by RT-qPCR. Each mouse is represented by a gray point showing relative values of parasite liver load normalized with the mean value from the ‘gp120’ control group. The box plot displays the upper and lower quartiles, median value and whiskers representing the data range. Dotted line denotes the limit of detection, defined as the mean signal from the no-template control.

Recognition of recombinant and sporozoite-derived antigens by vaccine-elicited antibodies was assessed using plasma samples collected prior to the mosquito-bite challenge (on day 21; i.e., one week post-boost). Plasma from each animal exhibited strong recognition of their respective recombinant immunogens by ELISA at a dilution of 1:50, with the exception of the animals immunized using the gp120 control protein (which showed poor immunogenicity, based on these data), suggesting that all recombinant PE immunogens elicited a robust antibody response (Fig. 1C). We next tested the recognition of natively expressed antigens by performing ELISA with plated whole wild-type *Py* sporozoites to assess the antigenic similarity between recombinant and natively expressed antigens. Our results revealed varying degrees of antibody reactivity (Fig. 1D) with respect to the background signal defined by the gp120 control condition (this protein is not expressed by the sporozoites), showing strongest reactivity in PyCSP- and PyTRAP-immunized plasma; moderate reactivity in plasma from PyCelTOS, PyP36 and PyP52 immunizations; and lower reactivity in plasma from the PyGAMA, PyHSP70-2, PyTRSP and PySSP3 immunizations. Despite the differences in the magnitude of signal in whole-sporozoite recognition, these results indicated that the antibodies elicited by the PE immunogens could recognize both recombinant and sporozoite-associated antigens, indicating a degree of antigenic similarity between recombinant and natively expressed antigens.

Following the immunization, mice were challenged with bites from 15 *Py*-infected mosquitoes (on day 28), and the parasite liver stage infection biomass was assessed by *Py*-specific 18S RT-qPCR relative to the control group immunized with the HIV-1 Env gp120 subunit. Mice immunized with single PE antigens exhibited a varying degree of protection from infection, manifested in a reduction in the parasite liver stage burden among the groups after challenge. The mice immunized with PyCSP or PyTRAP showed the most inhibition of infection, showing the lowest liver load, below the detection limit of the *Py* 18S RT-qPCR assay, representing >1000-fold reduction compared to the ‘gp120’ control (Fig. 1E). No other single-antigen vaccination regimen induced responses that reduced liver infection below the detection threshold. However, the remaining groups demonstrated reduction in parasite liver load ranging from 1.5- to 4.8-fold compared to the control group, indicating that liver infection was blocked to varying degrees. These results suggest that immunization with PyCSP or PyTRAP induces strong blocking antibodies against *Py* sporozoite infection, whereas the rest of the *Py* PE antigens elicited antibodies that were less inhibitory.

Notably, when accounting for the decrease in liver stage burden (Fig 1E), our plasma reactivity data imply that immunization groups with the highest signal against whole *Py* sporozoites (PyCSP and TRAP) also demonstrate the greatest protective effect against sporozoite infection. This may reflect differences in relative abundance (25) and/or accessibility of the target antigens at the sporozoite stage, or be due to nuances in their biological function and susceptibility to antibody inhibition. Regardless, these collective findings indicate that immunization with recombinant PE antigens induced strong antibody titers that recognize their cognate sporozoite-associated protein and block liver infection, albeit to varying degrees. However, the inhibition of sporozoite infection achieved by these PE immunogens (with the exception TRAP) is moderate compared to CSP-mediated inhibition, indicating that they are likely insufficient as stand-alone vaccines.

### Combined immunization of PE antigens with PyCSP identifies protective and non-protective co-immunogens

We next examined whether any of the PE immunogens could enhance PyCSP-mediated sterile protection in co-immunization/challenge experiments. Our standard immunization regimen to study sterile protection uses three doses of 20 μg of PyCSP in BALB/cJ mice and induces a strong protective effect during the mosquito-bite challenge (Suppl. Fig. 1A), which is too potent to detect enhancement of sterile protection by non-CSP antibodies. Thus, we evaluated several PyCSP immunization regimens to identify conditions resulting in sterile protection of 10–50% of mice per group, which would allow us to detect even nuanced statistically significant improvements in protection due to the addition of PE immunogens. In a pilot experiment, we observed that a 2-dose regimen with 5 μg of PyCSP induces 20% sterile protection (Suppl. Fig. 1B). Therefore, we chose the regimen consisting of 7 μg PyCSP combined with 13 μg PE immunogens or the control gp120 antigen (for a 20-μg total protein dose per injection in 20% Adjuplex) as our standard co-immunization strategy to assess changes in vaccine efficacy. Mice were immunized twice with combinations containing each single PE immunogen (or gp120 control) with PyCSP or gp120-only control, two weeks apart and then challenged with bites from 15 *Py*-infected mosquitoes 14 days post-boost (Fig. 2A). Using this regimen, the mice immunized with gp120 + PyCSP (n=30) showed 20% sterile protection (Fig. 2B), compared to the gp120 alone group (no PyCSP, n=30) showing 0% sterile protection (*p*=0.013, Barnard’s exact test; Table 2). Importantly, we observed that these control groups produced consistent outcomes over multiple experiments (Suppl. Table 1).

**Figure 2.**
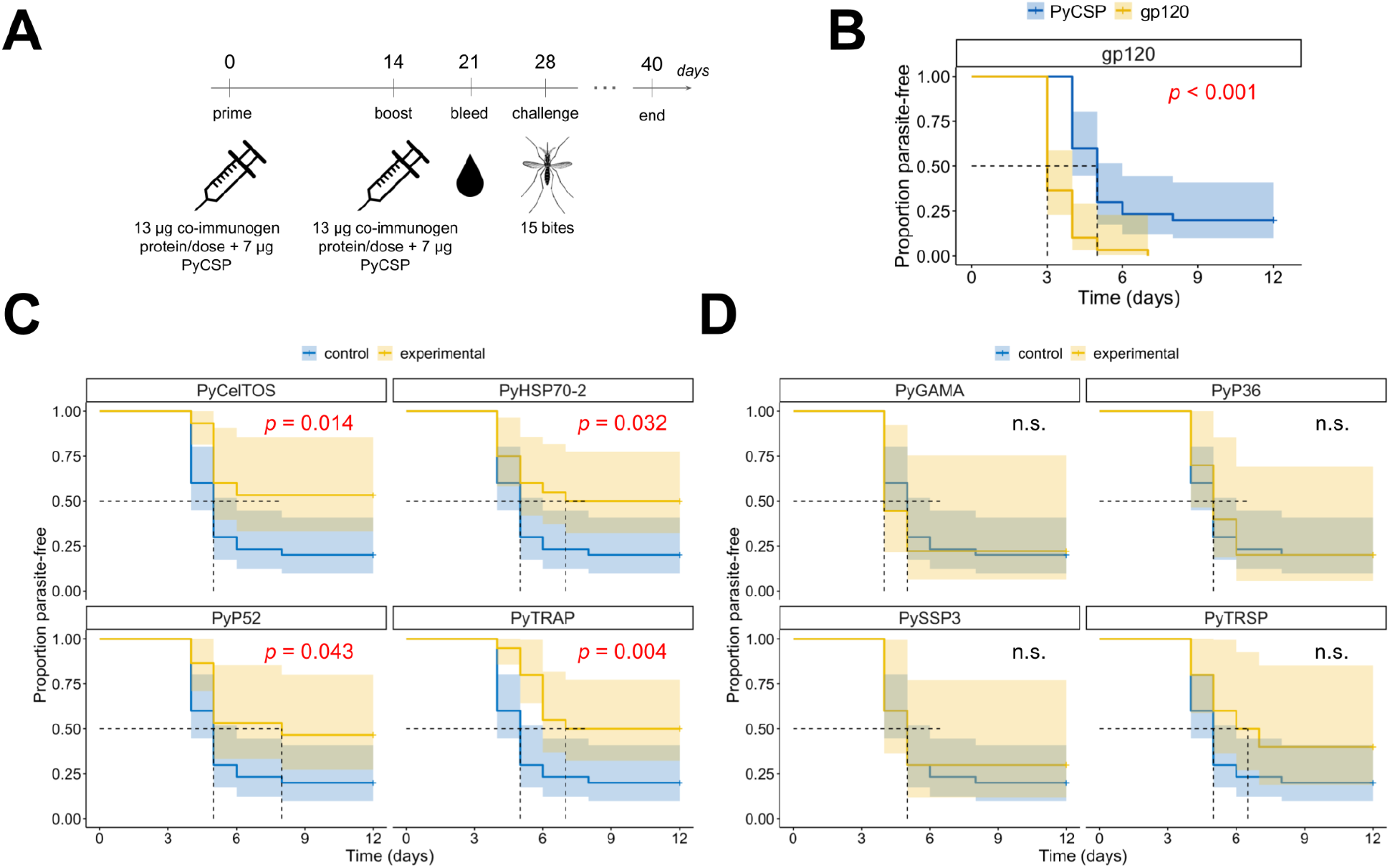
Co-immunization of PyCSP with recombinant sporozoite proteins delineates the protective and non-protective co-immunogens. **(A)** Groups of mice were immunized with a mix of 7 μg PyCSP and 13 μg of another PE antigen (as listed in Table 2 and Suppl. Table 1) following the same immunization-challenge strategy described in Fig. 1B. The level of sterile protection was determined by screening Giemsa-stained blood smears for parasitemia on days 3–12 post-challenge. **(B)** The ‘gp120+PyCSP’ group (blue line, n=30) showed the induction of significant sterile protection (20%), compared to the ‘gp120’ control group (yellow line, n=30, 0%), establishing the baseline of protection induced by PyCSP. **(C)** Co-immunizations with PyTRAP (n=20), PyHSP70-2 (n=15), PyCelTOS (n=20) and PyP52 (n=15), shown as a yellow line in each panel, demonstrate enhanced protection over ‘gp120+PyCSP’ group (blue line, same data as in **B**). **(D)** Co-immunizations with PyTRSP (n=10), PyGAMA (n=9), PyP36 (n=10) and PySSP3 (n=10), shown as a yellow line in each panel, were not statistically distinguishable from the “gp120+PyCSP” control group (blue line). The log-rank test was used to assess statistical significance of difference in each Kaplan-Meier plot, with *p*-values above 0.05 represented by ‘n.s.’

**Table 2.** Challenge outcomes following co-immunization (Barnard’s exact test)

| group | immunogen 1<br>(13 µg) | immunogen 2<br>(7 µg) | infected | total | protection<br>ratio (%) | *p*-value vs.<br>group A<sup>a</sup> | *p*-value vs.<br>group B<sup>a</sup> |
| --- | --- | --- | --- | --- | --- | --- | --- |
| A | *gp120* | n/a | 30 | 30 | 0% | *n/a* | *n/a* |
| B | *gp120* | PyCSP | 24 | 30 | 20% | **0.013** | *n/a* |
| C | PyHSP70-2 | PyCSP | 10 | 20 | 50% | *n/a* | **0.027** |
| D | PyTRAP | PyCSP | 10 | 20 | 50% | *n/a* | **0.027** |
| E | PyCeITOS | PyCSP | 7 | 15 | 53% | *n/a* | **0.026** |
| F | PyP52 | PyCSP | 8 | 15 | 47% | *n/a* | 0.075 |
| G | PyP36 | PyCSP | 8 | 10 | 20% | *n/a* | 1.000 |
| H | PyGAMA | PyCSP | 7 | 9 | 22% | *n/a* | 0.923 |
| I | PySSP3 | PyCSP | 7 | 10 | 30% | *n/a* | 0.585 |
| J | PyTRSP | PyCSP | 6 | 10 | 40% | *n/a* | 0.239 |

<sup>a</sup> Unadjusted Barnard's exact test *p*-value.

Among the PE antigen + PyCSP combinations tested, we identified three groups that showed significantly enhanced sterile protection over control: PyCelTOS (53%, *p*=0.026), PyHSP70-2 (50%, *p*=0.027) and PyTRAP (50%, *p*=0.027). Co-immunization with PyP52 resulted in 47% sterile protection, which was not significant, (*p*=0.075) (Table 2). When data were analyzed to account for the delay to onset of blood stage patency among the groups using the log-rank test, PyCelTOS (*p*=0.014), PyHSP70-2 (*p*=0.032) and PyTRAP (*p*=0.004) again showed statistical significance, however here PyP52 (*p*=0.043) also showed statistically significant improvement over the control gp120 + PyCSP group (Fig. 2C). The remaining PE antigen + PyCSP groups yielded lower proportions of sterilely protected animals following the challenge and were not otherwise statistically distinguishable from the control group (Table 2, Fig. 2D). Thus, of the 8 PE antigens tested, four improved sterile efficacy over the PyCSP immunization condition: PyCelTOS, PyTRAP, PyHSP70-2 and PyP52.

To quantitate the relative antibody response against each co-immunogen, and whether PE antigen titers associated with the occurrence of sterile protection, we assessed the total anti-PE-antigen immunoglobulin levels in each of the co-immunization groups (Fig. 3, Suppl. Fig. 2). For the majority of PE antigens tested, each of the animals immunized had half-maximal effective concentration (EC50) titers above 1:1,000, with PyTRSP yielding the highest EC50 titers (1:10,000–1:70,000). Notably, PyP36 and PyP52 had significantly lower EC50 titers with multiple animals measuring below the limit of detection (Figs. 3F, 3G). In addition, most mice immunized with the gp120 control protein had EC50 titers below the limit of detection (Figs. 3A, 3B), indicating poor immunogenicity of gp120 in these mice. Importantly, our data showed that regardless of the co-immunogen used, PyCSP elicited comparable EC50 titers across groups (1:1,000–1:45,000), demonstrating the independence of antibody responses to PyCSP from those elicited by PE immunogens under the co-immunization conditions used here. Correlation analysis between the EC50 titer and the protection status for each co-immunization group revealed no statistically significant correlations for either anti-CSP (Suppl. Fig. 3A) or anti-PE antigen response used in each respective group (Suppl. Fig. 3B). Likewise, no statistically significant correlation was detected between antibody EC50 titer and the day of blood stage parasitemia detection (Suppl. Figs. 4A, 4B). Taken together, our data indicate that the magnitude of antibody response alone is an insufficient predictor of either sterile protection status or of a delay to blood-stage patency.

**Figure 3.**
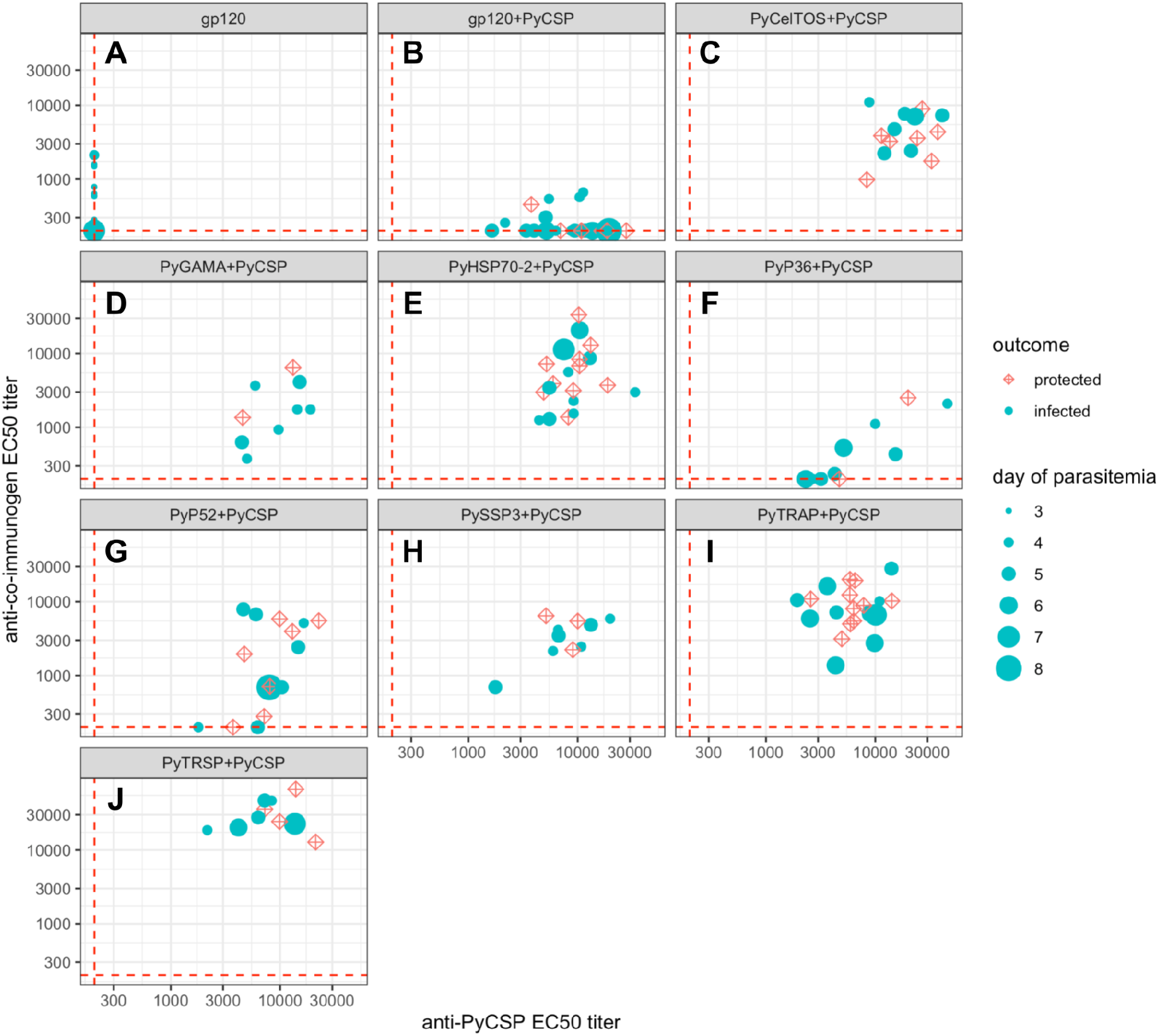
Co-immunization of PyCSP with each of the candidate proteins leads to robust antibody responses. Total antibody titers represented by the EC50 for each group were determined by ELISA using the “Day-21” plasma samples and plotted as the anti-PyCSP titer on x-axes and the anti-co-immunogen titer on y-axes. Each data point represents an individual mouse that either became infected (shown as the cyan filled-circle; size of the circle suggests the day when parasitemia was first detected) or remained free of detectable parasites (represented by a red crossed-square symbol). The EC50 limit of detection for all the antigens was set at 1:200 dilution (dashed red lines) and all the values below that threshold were adjusted to this value.

## Discussion

The RTS,S vaccine experience suggests that increasing the protective efficacy of CSP-based PE malaria vaccines should be a major priority for the field as it enhances the value of such a vaccine for malaria eradication efforts. However, developing a CSP-only malaria vaccine with high efficacy and durability of protection has been hampered by various roadblocks; and identifying additional PE vaccine antigens to augment anti-sporozoite immunity may be a promising strategy to fortify CSP-based vaccines. In our recent work, we have shown that plasma from mice immunized with recombinant TRAP—an exemplar PE antigen—inhibited sporozoites, and that passively transferred monoclonal antibodies could enhance anti-CSP-mediated protection from mosquito-bite challenge (23). Others have shown that active immunization with TRAP-CSP fusion proteins had protective effects in a similar experiment using the *Plasmodium berghei*-infected mosquito-bite challenge, although the individual contributions of anti-TRAP and anti-CSP responses were not completely defined (26). Additionally, a broader screen of co-immunogens derived from antigens expressed during liver stage development identified candidates that may be useful for eliciting T-cell responses (18). Here we show that several recombinant PE antigens normally expressed in sporozoites, when used in two-protein co-immunizations alongside PyCSP, can enhance sterile protection afforded by PyCSP alone.

Only four of the eight PE co-immunogens tested showed a statistically significant enhancement of anti-PyCSP response-associated sterilizing protection, even though all eight candidates elicited strong binding antibody responses. These differences in protective outcomes may be explained by the cellular localization differences among these protein targets, or by the degree to which their respective biological functions can be disrupted by vaccine-elicited interventions. In particular, CSP, TRAP, TRSP, P52, P36, CelTOS have been previously associated with invasion functions of the parasite (27–32). Previous work has also shown that immune response to CelTOS is associated with sterile protection from Plasmodium berghei challenge (33). HSP70-2 is an ER-resident chaperone that has been shown to associate with P36 and P52 (32). In contrast, neither GAMA nor SSP3 to date have been ascribed a role in PE stages. Nevertheless, in the context of vaccine targeting, disruption of protein function may not be required for protection, since antibody engagement of surface-exposed molecules alone may disrupt the ability of the parasite to traverse tissues (19) or mark the parasite for immune-mediated clearance (34).

Overall, our data show that the total polyclonal binding antibody levels resulting from immunization were an insufficient predictor of sterile protection in this study. It is possible that protection is associated with specific functional antibody subsets within the polyclonal antibody mixture either based on on-target epitope specificity (35), antibody isotype (10, 11) or off-target cross-reactivity (36). We and others had previously shown that the character of the polyclonal antibody response can be diverse enough to include antibodies capable of inhibiting the parasite as well as those that don’t have a detectable impact on the course of infection despite their ability to bind the antigen (23, 37–41). If this is the case here, then a more careful antibody response characterization with a focus on specific “vulnerable” epitopes would be required to describe the antibody response patterns that lead to protection. Additionally, this lack of correlation between antibody titers and protection status may be indicative of a more prominent role for cellular immunity, as has been observed in other similar systems (9, 42–44).

Our antibody binding data from the whole-sporozoite ELISA experiments imply that gross antigenic differences between the recombinant and sporozoite-associated proteins were unlikely to be the primary reason for lack of significant protection in single-protein immunizations. Despite the differences in glycosylation patterns of *Plasmodium* proteins (45), as compared with the mammalian recombinant versions, our data showed that the elicited antibodies recognized both the recombinant immunogens, as well as the native sporozoite-derived proteins. The variable signal strength in the whole-sporozoite ELISA experiments is likely indicative of different expression levels for each of these antigens in sporozoites (25), not of the strength of antibody response (the robustness of which is strongly indicated by the robust recognition of the recombinant versions). However, we cannot rule out the possibility that a subpopulation of antibodies may poorly recognize the native antigen, reducing the effective concentration of on-target circulating antibodies. A more careful dissection of epitope targeting and the identification of neutralizing epitopes in these PE antigen targets will provide insight into this, and would potentially enable the improvement of efficacy mediated by those antigens through iterative vaccine design.

To date, co-formulation with RTS,S in a clinical setting against *P. falciparum* infection has only been tested using TRAP recombinant protein, and in that study the RTS,S/TRAP/AS02 combination failed to protect against CHMI (46) despite prior work showing protection for RTS,S/AS02 alone (47). This created the concern that PE subunit vaccine co-immunization may be detrimental to CSP-mediated vaccine protection. In contrast, mixed regimens of RTS,S/AS01B and a viral vector-encoded multiple-epitope TRAP (ME-TRAP) construct elicited significant sterile protection from mosquito-bite challenge, albeit with no significant difference from the control RTS,S/AS01B-alone group (48, 49). Our mouse challenge data suggest that, when used in single-antigen immunizations, PyTRAP resulted in blocking of liver infection comparable to PyCSP, with all other immunogens in our panel yielding weak blocking activity. When used in combination with PyCSP, PyTRAP co-immunizations consistently resulted in enhanced probability of sterile protection compared to PyCSP-alone immunizations. Taken together with our recent finding that anti-TRAP mAbs can increase chances of sterile protection afforded by anti-CSP mAbs in a humanized mouse system (23), these data suggest that the use of recombinant TRAP in co-immunization strategies still holds promise and should be revisited.

In summary, our study provides an early stage proof-of-concept that immunization with recombinant CSP can be further enhanced by the addition of other sporozoite antigens, resulting in greater chances of sterile protection. All of the co-immunogens examined here have orthologs in *P. falciparum*. However, since our observations were made in a rodent malaria model, it is important moving forward to evaluate the efficacy of the *P. falciparum* versions in a physiologically relevant experimental system like the humanized-liver FRG-huHep mice (50). Some discrepancies have been observed previously between the rodent and human malaria models in vaccine development, and so confirmation of these findings against human-infective malaria is a critical next step. Additionally, it is likely that each antigen combination requires extensive optimization of dosages and formulations to enhance the relative antibody responses to achieve maximal efficacy, and careful evaluation of relevant epitope targets may allow rational design approaches. Future aspects of this study will also focus on testing the combinations of more than two antigens in achieving higher levels of sterile protection, and determining the optimal combinations and formulations to advance into the clinical setting. Overall, our findings pave the way for the development of combinatorial immunization regimens that may significantly enhance the protective properties of RTS,S and the next-generation subunit vaccines.

## Materials and Methods

### Constructs and protein production

Sequences encoding the ectodomains of *P. yoelii* proteins (Table 1) were codon-optimized and cloned into the pcDNA-3.4 expression vector flanked by the tissue plasminogen activator (tPA) signal sequence on the 5’ end, and the 8×His and AviTag sequences on the 3’ end. The two N-glycosylation sites present (identified by the NxS/T motif) in PyCSP were mutated to N×A (S27A,T348A), since it is currently unclear whether *Plasmodium* spp. carry out this protein post-translational modification (51).

Recombinant proteins were produced using the suspension HEK293-based expression system, as previously reported (52). Briefly, cultures of FreeStyle 293 (Thermo) cells were transiently transfected with the constructs using PEI MAX (Polysciences) at a 1:4 (w/w) DNA:PEI ratio and maintained for 5 days prior to harvesting the supernatants. Purification consisted of immobilized metal-affinity chromatography using HisPur resin (ThermoFisher) followed by gel filtration on a calibrated HiLoad Superdex 200pg column (Cytiva). Protein was then concentrated and stored at 4°C or flash-frozen (and stored at −20°C) until use. The HIV-1 Env gp120 control protein was produced, as previously described (53).

### Animal studies ethics statement

All procedures involving animals were performed in adherence to protocols reviewed and approved by the Institutional Animal Care and Use Committee (IACUC) at the Seattle Children’s Research Institute (Protocol ID# IACUC00487).

### Immunization

BALB/cJ mice (purchased from Jackson Laboratories, Bar Harbor, ME) were immunized twice—two weeks apart—with 20-μg total protein doses containing 13 μg of the candidate immunogens or control (gp120) and 7 μg of PyCSP, or 20 μg control protein in 20% Adjuplex adjuvant (Empirion LLC, Columbus, OH). Blood samples from immunized animals were collected on day 21 (i.e., 1 week post last immunization) through submandibular bleeding. The plasma was harvested by centrifuging blood samples at 4500×g for 10 min at 4°C and then immediately stored at −20°C until further use for the assessment of antibody responses.

The experiments were performed in two stages. First, a pilot stage with 5 mice per immunogen group was used to approximate protection outcomes, which was used to calculate the minimal numbers of animals per group needed to reach statistical power, according to Barnard’s exact test results (alpha=0.05, beta=0.2). In the second stage, several replicate experiments were performed to increase the group sizes and to capture the diversity of challenge efficiencies typically observed for multiple individual experiments.

### Mosquito-bite challenge and protection

The mosquito-bite challenge was done, as described previously (54). Briefly, the *Anopheles stephensi* mosquitoes were fed with *P. yoelii* (*Py*) 17XNL strain-infected blood meal and the infection rate was determined on day 9 post blood meal by analyzing mosquito midguts for the presence of oocyst. A small proportion of mosquitoes were dissected on day 14 post mosquito infection to verify the presence of sporozoites in the salivary glands. On day 15, immunized mice were anesthetized with Ketamine (100 mg/kg body weight)/Xylazine (10 mg/kg body weight) solution and placed above small cages each containing 15 mosquitoes, one mouse per cage. The mice were rotated for 10 min between the cages after every 50 seconds of incubation in order to minimize the variations in mouse infection and to maximize the probing events, as opposed to feeding. Sterile protection was determined by screening Giemsa stained thin blood smears, made from mouse tail snip bleeding, between days 3–12 post mosquito-bite challenge. Mice were considered protected if blood stage parasites were absent while examining at least 30 microscopic fields per mouse.

### Quantitation of antibody responses

Total antibody responses were quantitated by direct-immobilization ELISA. Antigens were diluted in 0.1M sodium bicarbonate buffer and incubated overnight at room temperature in wells of Immulon 2HB 96-well plates (Thermo Scientific) at a dose of 50 ng/well. Plates were then washed five times with PBS containing 0.02% Tween-20, which was performed between each step in the assay. Plates were then blocked against non-specific binding with 10% nonfat milk and 0.3% Tween-20 diluted in PBS for 1 hour at 37°C. Plasma samples obtained following immunization were heat-inactivated at 56°C for 30 minutes and diluted in PBS with 10% nonfat milk and 0.03% Tween-20 for a range of 1:20 to 1:5,598,720. Following an incubation for 1 hour at 37°C, the plates were washed and incubated in a 1:2000 dilution of HRP Goat Anti-Mouse Ig (BD, cat. #554002) in PBS with 10% nonfat milk and 0.03% Tween-20. Plates were then incubated for 1 hour at 37°C. Plates were developed using 50 μL/well of SureBlue Reserve TMB reagent (SeraCare Life Sciences Inc, cat. #5120-0083) and stopped after 3 minutes at room temperature by the addition of 50 μL/well of 1N sulfuric acid. Absorbance readings at 450 nm were performed using an ELx800 microplate reader (BioTek), and raw data were used to generate titration curves with the R package drc (version 3.0-1) to estimate the EC50 dilution values (sample data and fits shown in Suppl. Fig. 5).

### Antibody recognition of sporozoites in ELISA

The *Py* 17XNL sporozoites were harvested from the salivary glands of infected *Anopheles stephensi* mosquitoes as described previously on day 15 post infected blood meal (55). The sporozoites were diluted in phosphate-buffered saline (PBS; Corning cat. #21-040-CV) and deposited at 20,000 sporozoites/well in Immulon 2HB 96-well plates (Thermo Scientific), followed by an incubation overnight at 4°C. Parasites were then fixed by air-drying by aspirating all liquid from each well and incubating at room temperature for 25 minutes. Plates were then stored at −20°C until use. Following thawing, plates were blocked with 50 μL of 1% BSA (VWR, cat. #97061-422) in PBS for 1 hours at room temperature. Plasma samples obtained following immunization were heat inactivated at 56°C for 30 minutes and diluted in PBS with 1% BSA to achieve a 1:25 dilution. 50 μL of diluted sample was then added to the 50 μL block buffer on the plate in duplicate to achieve a final dilution of 1:50 and incubated at room temperature for 2 hours. Plates washed with 300 μL of PBS (pH 7.4) six times. Following washing, a 1:2000 dilution of HRP Goat Anti-Mouse Ig (BD, cat. #554002) diluted in PBS with 1% BSA was added to the plate and incubated for 1 hour at room temperature. Plates were again washed with PBS as described earlier. Plates were developed using 50 μL/well of SureBlue Reserve TMB reagent (SeraCare Life Sciences Inc, cat. #5120-0083) and stopped after 3 minutes at room temperature by the addition of 50 μL/well of 1N sulfuric acid. Absorbance readings at 450 nm were performed using an ELx800 microplate reader (BioTek).

### Determination of parasite liver load

The parasite load in the liver was determined as described previously (56). Briefly, the livers were perfused 45 hr post mosquito-bite challenge with 1× PBS and total RNA was extracted from the lower right caudate process of the caudate lobe of the liver from each mouse using miRNeasy kit (Qiagen: 217004). The complementary DNA (cDNA) was synthesized from 1 μg of total RNA using QuantiTect Reverse Transcription Kit (Qiagen: 205311) and quantitative PCR (qPCR) was performed using Bimake SYBR Green Master Mix (cat. #B21202) on QuantStudio 5 Real-Time PCR system. Briefly, 0.5 μL of diluted cDNA (1:10) was used in total 10 μL reaction volume to amplify *Py* 18S rRNA using 5’-GGGGATTGGTTTTGACGTTTT-3’ (forward primer) and 5’-AAGCATTAAATAAAGCGAATA-3’ (reverse primer), and the mouse *Gapdh* gene, as an internal control, using 5’-CCTCAACTACATGGTCTACAT-3’ (forward primer) and 5’-GCTCCTGGAAGATGGTGATG-3’ (reverse primer). The parasite liver load was calculated by comparative CT analysis and represented as 2^(-ΔΔCT)^ in relation to the gp120 control group (57).

The limit of detection was determined using a negative control reaction where no template cDNA was added.

### Data processing and statistics

Statistical tests were non-parametric (Barnard’s, log-rank) and employed with standard sidedness and 0.05 unadjusted *p*-value threshold. Due to small sample sizes, p-values were not adjusted for multiple comparisons. Raw data were processed using R (version 4.0.2) and packages tidyverse (version 1.3.1), drc (version 3.0-1), Exact (version 3.0), survival (version 3.2-13) and rstatix (version 0.7.0). Plots were generated using packages ggplot2 (version 3.3.5), ggpubr (version 0.4.0.999) and survminer (version 0.4.9).

## Data availability

All data are available in the article and supplementary information.

## Acknowledgements

We thank the staff of the vivarium at Seattle Children’s Research Institute for their support of the animal studies presented here. In addition, we thank Dr. Nana Minkah for his help with the study design and Tess Seltzer (of the insectary staff) for her diligent work in rearing the mosquitoes for these studies. This work was funded by NIH R01 AI117234 to DNS and SHIK.

## Author contributions

V.V. and H.P. contributed equally to this work.

Conceptualization and experimental design: V.V., H.P., S.H.I.K. and D.N.S.

Investigation: V.V., H.P., A.W., A.R., L.R., W.S., S.B.

Data analysis and visualization: V.V., H.P., P.T.E., S.H.I.K. and D.N.S.

Writing — Original draft: V.V.

Writing — Review and editing: V.V., H.P., P.T.E., S.H.I.K. and D.N.S.

Resources: S.H.I.K. and D.N.S.

Supervision, Project Administration and Funding Acquisition: V.V., S.H.I.K. and D.N.S.

## Competing Interests statement

The authors declare no competing interests.

## Notes

### Competing Interest Statement

The authors have declared no competing interest.

### Summary of Updates

Minor clarifications in the text were made and additional supplementary figures (Suppl. Figs. 2 and 5) were added to address comments during the review process.

