## Supplemental Information for "Co-immunization with pre-erythrocytic antigens alongside circumsporozoite protein can enhance sterile protection against *Plasmodium* sporozoite infection"

### Supplementary Information

#### Supplementary Tables

*Supplementary Table 1. Individual challenge experiment outcomes*

| group | condition | Experiment (protected/total) | | | | | | Total (protected/total) |
|---|---|---|---|---|---|---|---|---|
| | | *1* | *2* | *3* | *4* | *5* | *6* | |
| A | gp120 | 0/5 | 0/5 | 0/5 | 0/5 | 0/5 | 0/5 | 0/30 |
| B | gp120+PyCSP | 1/5 | 1/5 | 2/5 | 0/5 | 1/5 | 1/5 | 6/30 |
| C | PyHSP70-2+PyCSP | | 3/5 | | 4/10 | | 3/5 | 10/20 |
| D | PyTRAP+PyCSP | | 3/5 | | 4/10 | | 3/5 | 10/20 |
| E | PyCelTOS+PyCSP | 4/5 | | 2/5 | | 2/5 | | 8/15 |
| F | PyP52+PyCSP | | 2/5 | 3/5 | | 2/5 | | 7/15 |
| G | PyP36+PyCSP | | 1/5 | | | 1/5 | | 2/10 |
| H | PyGAMA+PyCSP | 0/5 | | | | | 2/4 | 2/9 |
| I | PySSP3+PyCSP | 2/5 | | | | 1/5 | | 3/10 |
| J | PyTRSP+PyCSP | 2/5 | | | | | 2/5 | 4/10 |

### Supplementary Figures

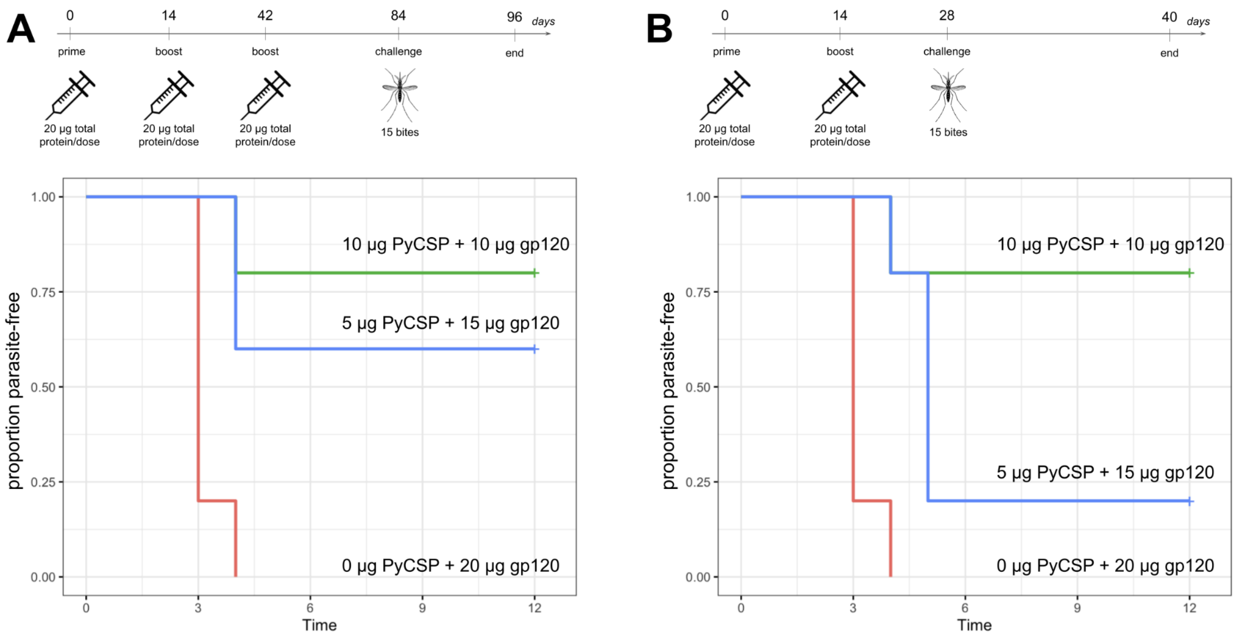


*Supplementary Figure 1. Testing the CSP dose and immunization numbers to achieve the sub-optimal sterile protection.*

Groups of mice (n=5) were immunized with the indicated amount of PyCSP per dose together with the gp120 control protein to achieve 20 µg of total protein content, injected in 20% Adjuplex. (**A**) shows a 3-dose regimen, followed by the challenge. (**B**) shows a 2-dose regimen, followed by the challenge.

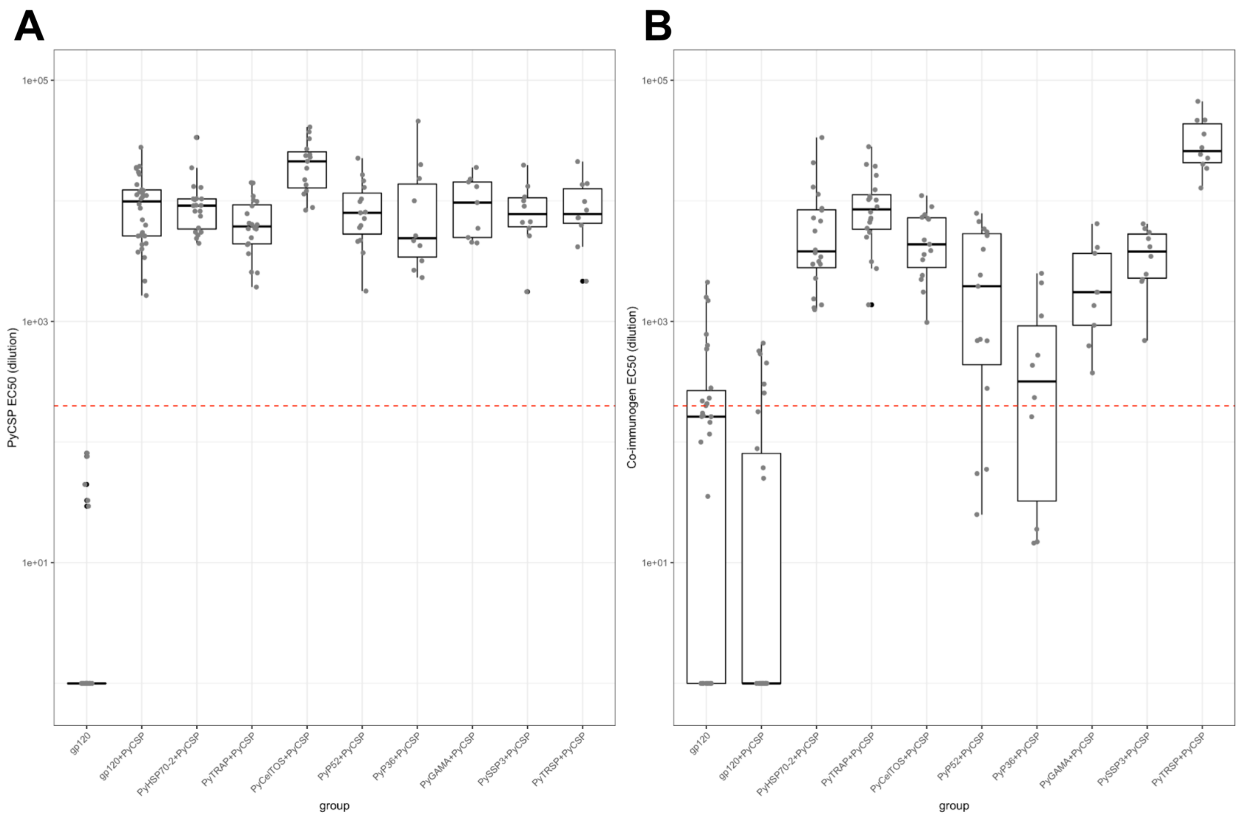


*Supplementary Figure 2. Total immunoglobulin titers for immunized mice (for Figure 3).*

EC50 values for PyCSP- (**A**) and co-immunogen-reactive immunoglobulins (**B**), shown in Figure 3, are plotted for each group. Each point represents a plasma sample; the conventional boxplot representation shows the per-group statistics including median, upper and lower quartile values. Red dashed line indicates the limit of detection (dilution of 1:200).

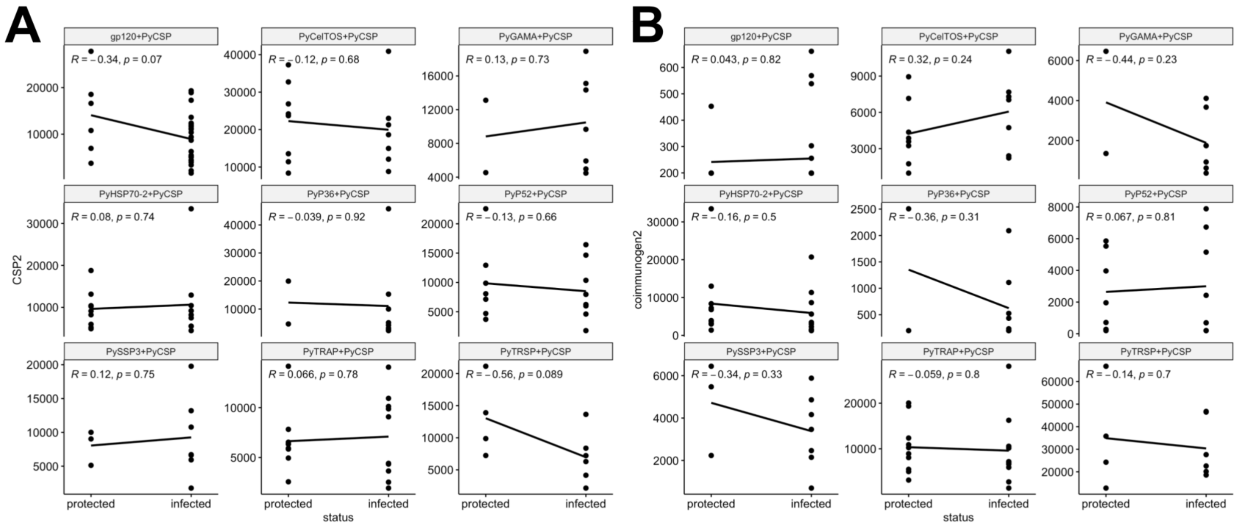


*Supplementary Figure 3. Correlation analysis of the antibody titers with the status of protection.*

Pearson correlation analysis against protection status using (**A**) the anti-PyCSP antibody titers (represented by EC50 values) or (**B**) the anti-co-immunogen antibody titers (represented by EC50 values).

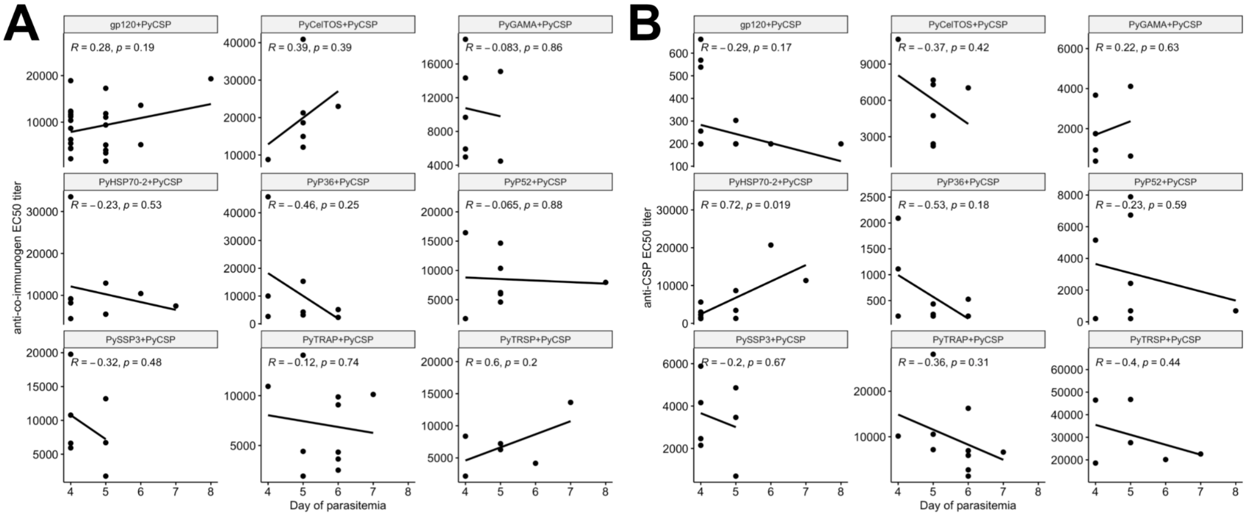


*Supplementary Figure 4. Correlation analysis of the antibody titers with the first day of detectable parasitemia.*

Pearson correlation analysis against first day of detectable parasitemia using (**A**) the anti-PyCSP antibody titers (represented by EC50 values) or (**B**) the anti-co-immunogen antibody titers (represented by EC50 values).

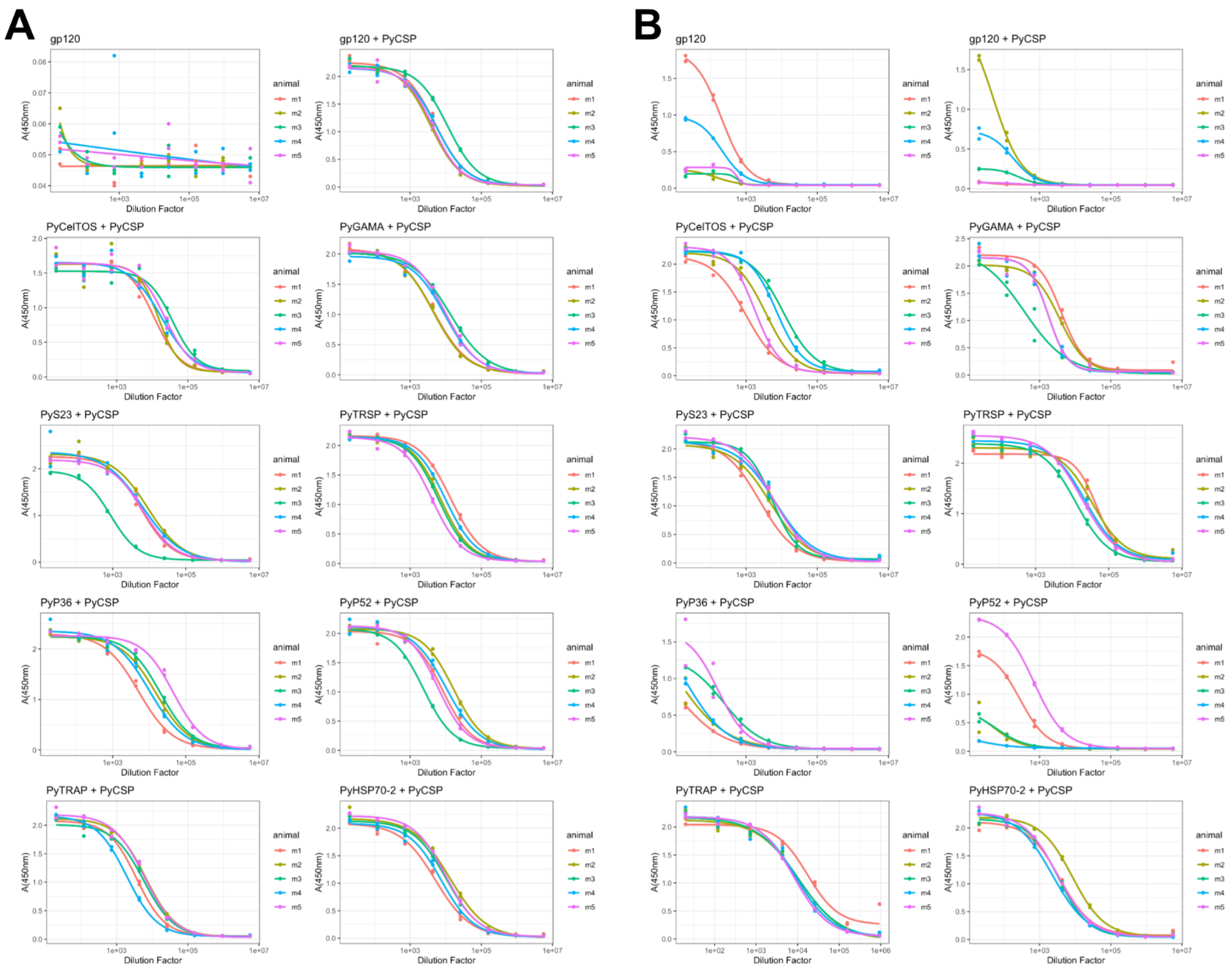


*Supplementary Figure 5. Sample immunized-mouse plasma titration curves used for estimating the EC50 values.*

Antigen-binding immunoglobulins in plasma samples from immunized mice were titrated using PyCSP (**A**) or the appropriate co-immunogen (**B**), measured in duplicate and shown as data points. Curve fits were constructed using the R package 'drc', shown as solid lines for each animal. Only a representative subset of data is shown here.
